# Resting-state Compensatory Remapping in Patients with Brain Tumour Before and After Surgery

**DOI:** 10.64898/2026.02.19.706837

**Authors:** Lucía Manso-Ortega, Ibai Diez, Lucía Amoruso, Garazi Bermudez, Santiago Gil Robles, Iñigo Pomposo, Manuel Carreiras, Ileana Quiñones

## Abstract

Brain tumours invade neural tissue, disrupting the functional organisation of neural networks. This disruption can trigger compensatory neuroplastic mechanisms that help preserve cognitive function despite pathological burden. Resting-state functional connectivity (rs-FC) provides a valuable approach for examining these alterations, yet the scope and trajectory of network-level changes remain unclear. In this study, we used rs-FC to characterise compensatory responses induced by left-hemisphere brain tumours affecting the integrity of the language network. The comparison of connectivity patterns at diagnosis and after surgery allowed us to track the temporal progression of these patterns. Relative to healthy participants, patients showed significant deviations in pre-surgical rs-FC, with widespread changes involving the dorsal attention, sensorimotor, frontoparietal, default mode, and cerebellar networks. These differences persisted post-surgically, with no significant longitudinal modifications. Network-level alterations also influenced topological properties: patients showed increased segregation while preserving global integration relative to controls. These properties changed significantly from pre- to postoperative period, with a postoperative increase in segregation and integration towards a pattern more closely resembling that of healthy controls. Furthermore, preoperative network segregation was identified as a predictor of postoperative cognitive recovery. These findings suggest that brain tumours induce network remapping, affecting functional connectivity far beyond the regions strictly related to the affected region. The observed changes in network topological organisation likely reflect the adaptive role of domain-general networks in facilitating homeostatic processes. Surgical strategies aimed at preserving these changes seem to promote sustained cognitive function, regardless of malignancy grade or lesion extent.

**Key points:**

- Presurgical patients show widespread inter- and intrahemispheric rs-FC alterations, reflecting tumour-driven network remapping.
- Network integration and segregation show longitudinal recovery, approaching neurotypical topology four months post-surgery.
- Preoperative network segregation predicts postoperative cognitive recovery.

**Importance of the Study:** Brain tumours disrupt large-scale brain networks, yet the extent, temporal dynamics, and cognitive relevance of these alterations remain poorly understood. Using resting-state functional connectivity, this study provides a longitudinal, network-level characterisation of tumour-induced neuroplasticity in patients with left-hemisphere lesions affecting language-related regions. Compared with prior work focused on local effects or single networks, our results demonstrate widespread inter- and intrahemispheric connectivity changes involving multiple domain-general resting-state networks, highlighting extensive network remapping beyond lesion boundaries. Importantly, we show that postoperative recovery is accompanied by a rebalancing of network integration and segregation toward a neurotypical configuration. Critically, preoperative network segregation emerges as a predictor of postoperative cognitive recovery, identifying a potential biomarker of resilience. These findings have direct translational relevance, suggesting that preserving and supporting large-scale network organisation during surgical planning may promote cognitive outcomes independently of tumour grade or lesion extent.

## 1. Introduction

Brain tumours pose a significant challenge to the integrity of human brain networks, yet they simultaneously trigger adaptive neuroplastic responses ^1–4^. In the current study, we aimed to track signs of adaptation in the global organisation of resting-state networks (RSNs) and their temporal evolution in a cohort of patients with left-hemisphere brain tumours, assessed both before and after surgical resection. In this pathological context, neuroplasticity is considered fundamental for preserving cognitive functions across the pre- and postoperative stages ^5^. A growing body of evidence suggests that resting-state functional connectivity (rs-FC), which reflects patterns of spontaneously synchronised neural activity, offers a powerful window into network-level adaptations ^6^. The use of rs-FC is a promising tool for assessing the organisation of large-scale brain systems in clinical populations, including patients with brain tumours ^7–10^.

The influence of brain tumours seems to extend beyond focal regions to affect global brain connectivity ^9^, affecting multiple RSNs. For instance, even when specific networks such as the language network are disrupted, compensatory mechanisms may emerge, including increased cerebro-cerebellar connectivity, potentially reflecting an adaptive response aimed at preserving functional performance ^1^. Further, the nature of changes in networks such as the Default Mode Network (DMN) or the Fronto-Parietal Network (FPN) is variable, with reports of both increases and decreases in connectivity ^1,11–13^. Critical network hubs appear particularly vulnerable ^14^, and significant topological deviations from healthy individuals have been documented, with altered patterns in RSNs distant from the tumour correlating with cognitive deficits ^8^. New metrics are being developed to quantify these alterations ^15^ and linking tumours’ biological features with RSNs disruptions ^16^, towards finding potential common underlying mechanisms.

Growing interest in functional plasticity in the context of brain tumours emphasises the importance of understanding how RSNs evolve after surgical resection. However, longitudinal studies remain scarce, and their findings are often hard to generalise, largely due to small sample sizes and the clinical heterogeneity of patient populations. Reviews by Cargnelutti, Ius, Skrap, et al. (2020) ^5^ and Sighinolfi, Mitolo, Testa, et al. (2022) ^6^ underscore this lack of empirical evidence, reporting only nine and five longitudinal studies, respectively, from which some are single-case reports. Within the available evidence, we find reports suggesting dynamic changes in interhemispheric connectivity ^17^, associations between the integrity of specific networks and post-operative cognitive performance ^18^, and the preservation of certain networks after surgery ^19,20^. Functional proximity to the tumour, rather than anatomical distance, seems to better predict network disruption ^21^. Overall, post-operative recovery is often accompanied by changes in activation patterns suggestive of active remapping ^6,22^. Recent findings, including our own, support these observations, showing that functional alterations are already present before tumour resection. These alterations include not only language network remapping ^23,24^ but also changes in the spatiotemporal dynamics of RSNs ^22^, manifested as reduced spontaneous neural activity in the cerebellum, diminished connectivity between DMN regions, and weakened interhemispheric connections between motor areas ^9^.

Given the distributed impact of brain tumours on functional communication within neural networks, recent research has increasingly relied on graph theory ^3,25,26^, which conceptualises the brain as a complex system of nodes (brain regions) and edges (functional or structural connections) ^27^. Graph-theoretical metrics are particularly informative in this context, as focal lesions can disrupt the topological organisation of brain networks in ways that simple measures of connectivity strength may fail to capture. This limitation is further compounded by substantial interindividual variability in the size and location of tumours, which reduces the sensitivity of conventional group-level analyses ^3,25,26^.

Recent studies have consistently reported alterations in the topological properties of RSNs in tumour patients relative to healthy controls, including changes in global efficiency, path length, and network segregation ^3,4,9,28^. To address these challenges, the present study focuses on characterising global network integration and segregation, both of which have emerged as sensitive predictors of neuroplasticity in pathological conditions ^9,25,26,28–30^. Global efficiency reflects the overall capacity of the brain network—despite tumour-induced disruption—to transmit information rapidly and reliably across distributed regions. In parallel, segregation quantifies the network’s ability to self-organise into functionally specialised modules ^31^. Together, these complementary metrics offer a comprehensive framework for understanding how information is integrated and partitioned within the tumour-affected brain.

In the current study, we investigate how neuroplasticity extends beyond domain-specific functions to influence the global organisation of RSNs over time. Specifically, we examine network-level alterations in rs-FC patterns in patients with left-hemisphere lesions, both before and after surgery, using a group of healthy participants as a reference. To characterise the evolution of these connectivity patterns in relation to cognition—and to assess whether they reflect compensatory processes—we combined two complementary analytical approaches: (i) a multivariate contrast of estimated ROI-to-ROI network changes, controlling for demographic and clinical factors, and (ii) graph theoretical analysis, focusing on global metrics of network integration and segregation to capture large-scale, tumour-related organisational remapping. Clinical variables (e.g., tumour type and size) and demographic data (e.g., age, sex) were included as moderators of the relationship between network alterations and cognitive trajectories. Overall, this integrative approach aims to characterise long-term connectivity changes induced by brain tumours, providing a step toward the development of personalised clinical strategies that incorporate resting-state neuroimaging markers into routine patient assessment and decision-making.

## 2. Methods

### 2.2. Participants

We included seventy-two patients with primary brain tumours without metastases, and a cohort of sixty-eight healthy participants used as a control group. All patients were evaluated between one and three weeks after the episode that prompted their first consultation with the Neurosurgery Department at Cruces University Hospital (Bilbao, Spain). Both patients and controls underwent a comprehensive protocol combining advanced functional and structural neuroimaging techniques with an extensive cognitive assessment spanning multiple domains. From the initial patient cohort, only those who meet the following inclusion criteria were retained for analysis: (1) a lesion in the left-hemisphere involving at least one critical language node, (2) FLAIR imaging consistent with diffuse low-grade glioma, without hyperintensities suggestive of high-grade pathology, (3) initial surgical treatment performed within four weeks of diagnosis, and (4) no postoperative cognitive intervention.

This resulted in thirty-five patients (mean age = 45.34 years; SD = 13.65), thirteen of whom were women. All patients had normal hearing and normal or corrected-to-normal vision. The project included a longitudinal follow-up of up to one year after surgery. However, due to the clinical complexity of oncology patients and variability in treatment, only a subset of twenty-four individuals completed both postoperative rs-fMRI and behavioural assessments, which were conducted approximately four months after surgery (mean interval between assessments = 132 days; SD = 48.47). A summary of tumour characteristics, including grade according to the WHO’s classification ^32^, is provided in Table 1, describing the histopathological analysis of tissue samples collected during surgery. As shown, some patients exhibited no high-grade features on imaging but displayed histological or genetic evidence of high-grade disease.

**Table 1.** Description of tumour characteristics (n = 35) based on the World Health Organisation (WHO) (Louis et al., 2021) pathophysiological classification.

| **Overall tumour classification** |  |
| --- | --- |
| High-grade glioma | 14 (40%) |
| Low-grade glioma | 18 (51.4%) |
| Other | 3 (8.1%) |
| **Tumour classification according to affected cell type** |  |
| Glioblastoma | 6 (17.1%) |
| Astrocytoma | 14 (40%) |
| Oligodendroglioma | 10 (27%) |
| Oligoastrocytoma | 1 (2.9%) |
| Xanthoastrocytoma | 1 (2.7%) |
| Dysembryoplastic | 2 (5.7%)^ |
| Cortical Dysplasia* | 1 (2.9%)^ |

All included tumours induce neuroplastic changes due to their impact on neural tissues, including the three with non-glial tumours (^). Extra-axial lesions were excluded from this study. \* Not considered in the WHO classification as it is not a neoplastic lesion.

A power estimation was conducted based on a repeated-measures design (two time points: pre- and postoperative) using G*Power (version 3.1.9.7) ^33^. Assuming a moderate within-subject correlation (*r* = 0.6) and a two-tailed significance level of α = 0.05, a sample size of twenty-four participants yields an estimated power (1–β) of between 0.80 and 0.96 to detect moderate to large effect sizes (Cohen’s d ≈ 0.6–0.8). This level of sensitivity is adequate to detect meaningful longitudinal changes in cognitive performance, particularly given the dimensionality reduction achieved through the use of composite measures, which will be described in the following sections. The group of healthy participants consisted of 68 individuals (25 males, 43 females; mean age, 45.18 years; SD, 13.29 years), all of whom had normal hearing and normal or corrected-to-normal vision, and no history of neurological or psychiatric disorders.

### 2.3. Ethics statement

Data collection was approved by the Ethics Board of the Euskadi Committee and the Ethics and Scientific Committee of the Basque Center on Cognition, Brain, and Language (BCBL) (protocol code PI2020022, date of approval: 26 May 2020, and protocol code: 270220SM, respectively). The study protocol was conducted in accordance with the Declaration of Helsinki for experiments involving humans. At the beginning of each session, all participants received instructions about the techniques and tasks to perform. Informed consent was acquired before data collection. Healthy participants were economically compensated for their time.

### 2.4. Cognitive assessment

All participants underwent a series of standardised neuropsychological and behavioural evaluations. For patients, cognitive assessments were conducted twice: approximately one week before surgery and again ∼four months post-intervention. This allowed for a direct comparison of their cognitive performance pre- and post-surgery. Healthy participants completed this evaluation only once.

Evaluations mainly assessed three domains: language, working memory, and executive functions. To quantify performance within each domain, we created composite scores. These were constructed by aggregating the standardised performance values from related tasks (as detailed in Table 2). Composite variables are a valuable tool, particularly in clinical settings, as they can effectively combine multiple measures ^34^, manage missing data (see Supplementary Material Table 1S for full details on data availability), and capture the multifaceted nature of cognitive abilities ^35^. The available behavioural individual scores were used to derive these proxies, with the mean calculation adjusted according to the number of contributing measures available for each individual. The number of non-missing values for all composite variables at each time point can be found in the Supplementary Material (Table 1S). Individual composite scores were used to generate a cognitive progress index by comparing postoperative and preoperative assessments, capturing longitudinal changes in language, working memory, and executive functions, respectively. To ensure comparability of differences across cognitive domains, these values were normalised by dividing by the standard deviation of each metric. As a result, the obtained scores were centred and standardised.

**Table 2.**
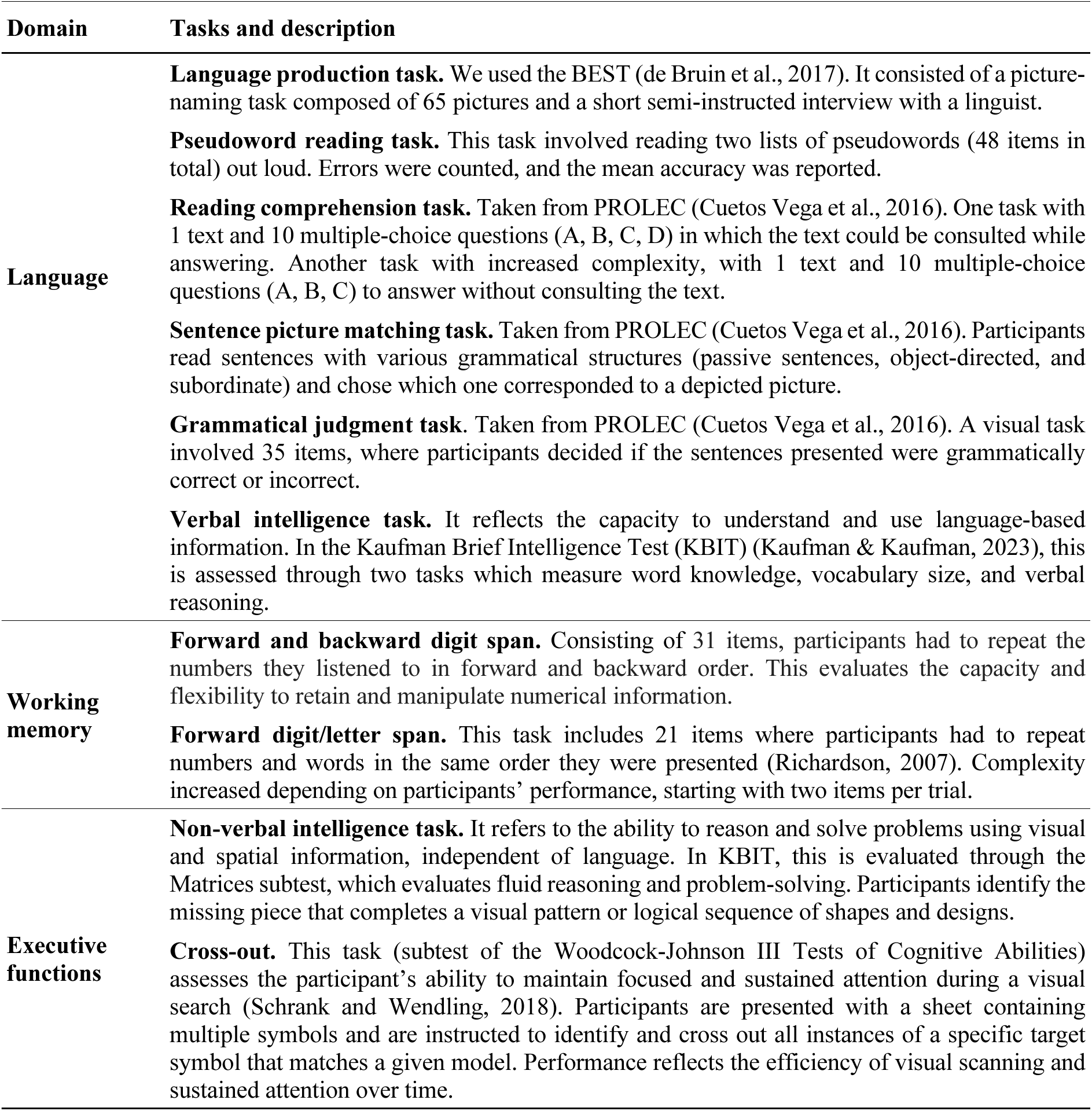
Cognitive assessment. This table provides a comprehensive overview of the tasks included in the assessment to generate the composite scores for each cognitive domain.

### 2.5. MRI Data Acquisition

All MRI data were acquired using a 3T Siemens Magnetom Prisma Fit scanner (Siemens AG, Erlangen, Germany). To obtain high-resolution images, a 3D ultra-fast gradient echo (MPRAGE) pulse sequence was employed, using a 64-channel head coil. The T1-weighted images were acquired with the following parameters: 176 contiguous sagittal slices, voxel resolution of 1x1x1 mm³, repetition time (TR) = 2530 ms, echo time (ET) = 2.36 ms, 256 image columns, 256 image rows, and a Flip angle (Flip) = 7°. Similarly, the T2-weighted images were acquired with 176 contiguous sagittal slices, voxel resolution of 1x1x1 mm³, RT = 3390 ms, ET = 389 ms, 204 Image columns, 256 Image rows, and Flip = 120°.

Resting-state functional MRI (rs-fMRI) data for patients were acquired during a period spanning approximately 10 years, and two different sequences were used. For Sequence A (19 patients), rs-fMRI data were acquired with a voxel resolution of 3mm³, a repetition time (TR) = 2000 ms, an echo time (TE) = 30 ms, and a flip angle (Flip) = 78°. Each volume comprised 32 slices acquired with a matrix size of 91 x 109, and had an echo spacing of 64 ms. A total of 180 echo planar images were collected. The image columns and rows were 384 and 384, respectively, and the acceleration factor was 0. For Sequence B (16 patients), rs-fMRI data were acquired with a voxel resolution of 2mm³, a TR = 850 ms, a TE = 35 ms, and a Flip = 56°. Each volume consisted of 66 slices acquired with a matrix size of 91 x 109, and had an echo spacing of 88 ms. A total of 1058 echo planar images were collected. The image columns and rows were 792 and 792, respectively, and the acceleration factor was 0.

To mitigate potential confounding effects arising from different acquisition parameters, the sequence type (A or B) was included as a covariate in all relevant statistical analyses. Healthy participants were all scanned with sequence B. Detailed parameters for each rs-fMRI sequence are presented as Supplementary Material Table 2S. During the rs-fMRI scan, which lasted approximately 10 minutes, participants were instructed to lie still with their eyes open, fixate on a central crosshair displayed on a screen, remain awake, and let their minds wander without focusing on any specific thought.

### 2.6. Lesion reconstruction

Lesions were reconstructed under the supervision of the neurosurgeons in charge of the patient’s surgical intervention (G.B.). Masks were manually drawn slice by slice on the native space using MRIcro-GL (Rorden & Brett, 2000). To identify the lesion-affected area, we relied on coregistered T1- and T2-weighted images and multiple intensity thresholds. These individual lesion masks were then normalised to the Montreal Neurological Institute (MNI) standard space, and the alignment between the reconstructed lesion and the lesion in the native space was checked. We created a volume of interest (VOI) for each patient at each time point and then smoothed it. This VOI was used to estimate the tumour volume (cm^3^) and the extent of resection. The extent of resection (cm^3^) was estimated on postoperative imaging as (Volume of preoperative 3D Tumour Reconstruction / postoperative resection) * 100/preoperative tumour volume. An overlap lesion map representing the spatial distribution of tumours across the 35 patients included in this study is depicted in Figure 1.

**Figure 1.**
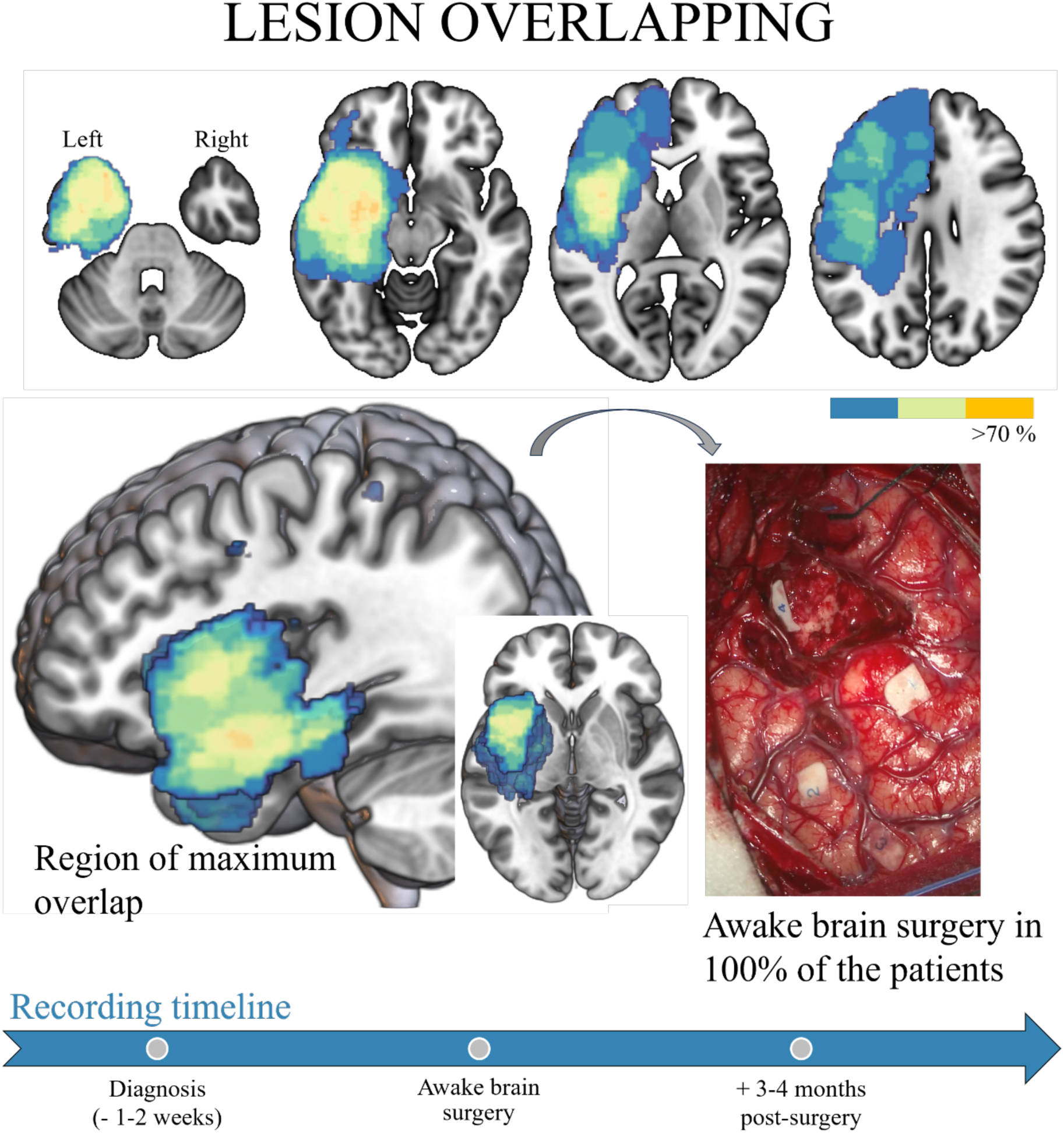
Lesion overlap map for the patient group. The colour bar indicates the percentage of overlapping tumours at each voxel, ranging from blue (low overlap) to orange (high overlap). The 3D reconstruction of the lesion was based on imaging studies performed one to three weeks before surgery, which coincided with the time of diagnosis in all patients. As depicted in the right-hand panel, the surgical procedure adhered to established protocols for awake craniotomy, incorporating intracortical electrical stimulation to delineate functional boundaries. The bottom arrow marks the longitudinal approach and the associated recording intervals.

### 2.7. MRI Data Preprocessing and Analysis

Functional resting-state data images were preprocessed using SPM12 (Welcome Department of Cognitive Neurology, London, UK) using in-house Matlab (2021b release, Mathworks, Inc, Natick, MA) codes. Raw functional scans were slice-timed corrected with the middle slice as the reference. After commissure adjustment, scans were realigned, unwarped, and coregistered with the anatomical T1 and normalised to MNI space using the unified normalisation segmentation procedure. Then, scans were smoothed with an 8mm³ isotropic Gaussian kernel. As part of the preprocessing, through CONN (RRID: SCR_009550) release 22.a ^36^, potential outliers scans were identified using artifact detection tools (ART) (2011 release, Cambridge, MA), as acquisitions with framewise displacement above 0.9 mm or global BOLD signal changes above 5 standard deviations (Power et al., 2014), and a reference BOLD image was computed for each subject by averaging all scans excluding outliers. In addition, functional data were denoised using a standard denoising pipeline ^37^, including the regression of potential confounding effects characterized by white matter time series (5 CompCor noise components), CSF time series (5 CompCor noise components), session effects and their first order derivatives (2 factors) ^38^, art regressors (106 components) to account for both outliers and movement and a constant term (1 factor) within each functional run, followed by bandpass frequency filtering of the BOLD time-series (Hallquist et al., 2013) between 0.008 Hz and 0.09 Hz.CompCor (Behzadi et al., 2007; Chai et al., 2012) noise components within white matter and CSF were estimated by computing the average BOLD signal as well as the largest principal components orthogonal to the BOLD average within each subject’s eroded segmentation masks. From the number of noise terms included in this denoising strategy, the effective degrees of freedom of the BOLD signal after denoising were estimated to range from 20.9 to 144.7 (average 125.1) across all subjects in the comparison between patients and healthy participants and 99.1 to 288.4 (average 196.6) in the longitudinal analysis.

As a first-level analysis, ROI-to-ROI connectivity (RRC) matrices were estimated, characterising the rs-FC between each pair of regions among 32 HPC-ICA network ROIs focused on the sensorimotor network, language network, visual network, executive control network, lateralized frontoparietal network, salience network, dorsal attention network, and default mode network (DMN) ^36^. FC strength was represented by Fisher-transformed bivariate correlation coefficients from a general linear model (weighted-GLM), estimated separately for each pair of ROIs ^37^, characterising the association between their BOLD signal time series. To compensate for possible transient magnetisation effects at the beginning of each run, individual scans were weighted by a step function convolved with SPM12 canonical hemodynamic response function and rectified.

### 2.8. Second-level analysis comparing patients and healthy participants

To compare patients before surgery and healthy participants, group-level analysis was performed using a General Linear Model (GLM). For each connection, a separate GLM was estimated, with first-level connectivity measures at this connection as dependent variables and group (patient or healthy participant) as an independent variable, with sequence and age as covariates (y ∼ 1 + Group + Sequence + Age) to evaluate the effect of group. Connection-level hypotheses were evaluated using multivariate parametric statistics with random effects across subjects and sample covariance estimation across multiple measurements. Inferences were performed at the level of individual networks (groups of related connections), as recommended to compare independent samples (Zalesky et al., 2012). Non-parametric statistics from Network-Based Statistics analysis were estimated (NBS) ^39^, with 1000 residual-randomisation iterations. Results were thresholded using a combination of a cluster-forming *p* < 0.001 connection-level threshold and a false discovery rate (*p*-FDR < 0.05) network-mass threshold ^40^.

### 2.9. Second-level analysis comparing patients before and after surgery

To assess rs-FC changes before and after surgery within the patient group (*n* = 24), a similar GLM approach was used, modelling within-subject changes. First-level rs-FC measures served as dependent variables. The model included time point (pre- and post-surgery) as a within-subject factor, with age and acquisition sequence as covariates (between-subjects model specification: y ∼ 1 + Sequence + Age (1 0 0); within-subjects model specification: y ∼ pre + post1 (−1 1)). Further, we tested the effect of tumour grade (high- vs. low-grade) on rs-FC and the relationship between rs-FC and progress in three cognitive variables: intelligence (including verbal and non-verbal abilities), language, and working memory. Inferences were performed at the level of individual networks. These variables were included as covariates in separate GLM analyses. Inferences for longitudinal analyses were also performed at the network level using NBS with the same *p* < 0.001 primary threshold and *p*-FDR < 0.05 network-level correction.

### 2.10. Graph-theoretical analysis of functional network reorganization

To obtain an index summarising the impact of the lesion on presurgical resting-state functional network organisation, two global graph-theoretical metrics were computed and used as predictors: global efficiency and functional segregation. Both measures were derived from weighted functional connectivity networks using the Brain Connectivity Toolbox (BCT ^41^) and correspond to the global efficiency index (*Eglob* in BCT) and the maximised modularity index (*Q* in BCT), respectively. Global efficiency was used as a measure of network integration and is defined as the average inverse shortest path length between all pairs of nodes in the network:

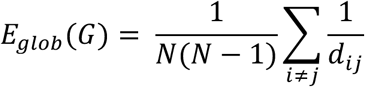

where 𝑑*_ij_* is the length of the shortest path between nodes 𝑖and 𝑗, computed on a matrix of edge lengths 𝐿 = derived from the weighted adjacency matrix as:

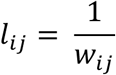

For weighted functional networks, connectivity weights were interpreted as connection strengths and converted to edge lengths by taking the inverse of the absolute weight values before shortest-path computation. In this way, both positive and negative correlations contributed to network topology through their absolute magnitude.

Functional segregation was quantified using the modularity quality index 𝑄, which captures the extent to which the network is organised into densely interconnected modules with sparse inter-modular connectivity. Modularity was defined as:

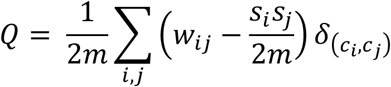

where 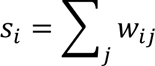 is the strength of node 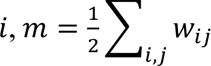 is the total weight of the network, 𝑐*_i_* denotes the community assignment of node 𝑖, 𝛿(𝑐*_i_*, 𝑐*_j_*) is the Kronecker delta, equal to 1 if 𝑐*_i_*, = 𝑐*_j_* and 0 otherwise. Higher 𝑄 values indicate stronger segregation and greater functional specialisation.

Both global efficiency and modularity have been widely applied in pathological contexts and have demonstrated high sensitivity to network-level alterations associated with increasing pathological burden in neurological conditions, including neurodegenerative diseases and epilepsy ^42–44^. Graph-theoretical measures provide a principled framework for characterising the impact of pathology on large-scale functional architecture while minimising dependence on individual nodes or edges and reducing data dimensionality. By jointly considering metrics of integration (global efficiency) and segregation (modularity), it is possible to estimate the balance between network-wide communication and functional specialisation ^45^. This integrated approach enables a robust assessment of lesion-related and surgery-induced changes in functional network organisation and their temporal progression, while reducing the degrees of freedom in the statistical model—an essential consideration for detecting small-to-moderate effects in a limited sample (n = 24).

### 2.11. Moderation analysis

A moderation analysis was conducted to examine whether the relationship between the connectomic measures and cognitive progress in language, working memory, and executive functions was mediated by clinical (i.e., malignancy tumour grade, and lesion volume) and demographic variables (i.e., age). The moderation analysis was performed using structural equation modelling (SEM) with maximum likelihood (ML) estimation ^46^, as implemented in JASP (version 0.95.1; JASP Team, Amsterdam, The Netherlands). Three potential moderators were included in the model: age (in years), malignancy tumour grade level (a four-level classification), and total lesion volume (in cm³). The effect of sex was controlled as a background confounder. Indirect, direct, and total effects were estimated for each path, and statistical significance was assessed using z-values with corresponding *p*-values. In addition, 95% confidence intervals were computed for all parameter estimates using bias-corrected bootstrap resampling with 1,000 iterations. Model fit was evaluated through χ² tests, path coefficients, and residual covariance analysis. The strength of associations between variables was further quantified with Pearson’s correlation coefficients, and effect sizes were expressed using Fisher’s z transformation. All reported values are based on two-tailed tests with α set at 0.05.

### 2.12. Sensitivity analyses

To examine potential effects derived from combining different fMRI acquisition sequences, we implemented a harmonisation strategy following the methodology approach discussed in ^47^. Individual time series were normalised, ensuring homogeneous dynamic ranges and consistent minimum and maximum values across scanners using two different approaches: Z-scores and ComBat, an empirical Bayesian method ^47–49^. Results obtained after harmonisation showed high concordance with those from the initial analyses, with no significant differences observed between groups (see Figure 2). In addition, we controlled for possible demographic effects, specifically age and sex, on the connectomic metrics derived from the fMRI data by including them as covariates in the group comparisons. After covariate adjustment, all reported group differences remained statistically significant. Additionally, we assessed the potential influence of data quality measures (e.g., motion and signal-to-noise ratio) by including them in the statistical analyses, which revealed no significant effects. These sensitivity analyses confirm the robustness and stability of the main findings against potential methodological or demographic sources of variability.

**Figure 2.**
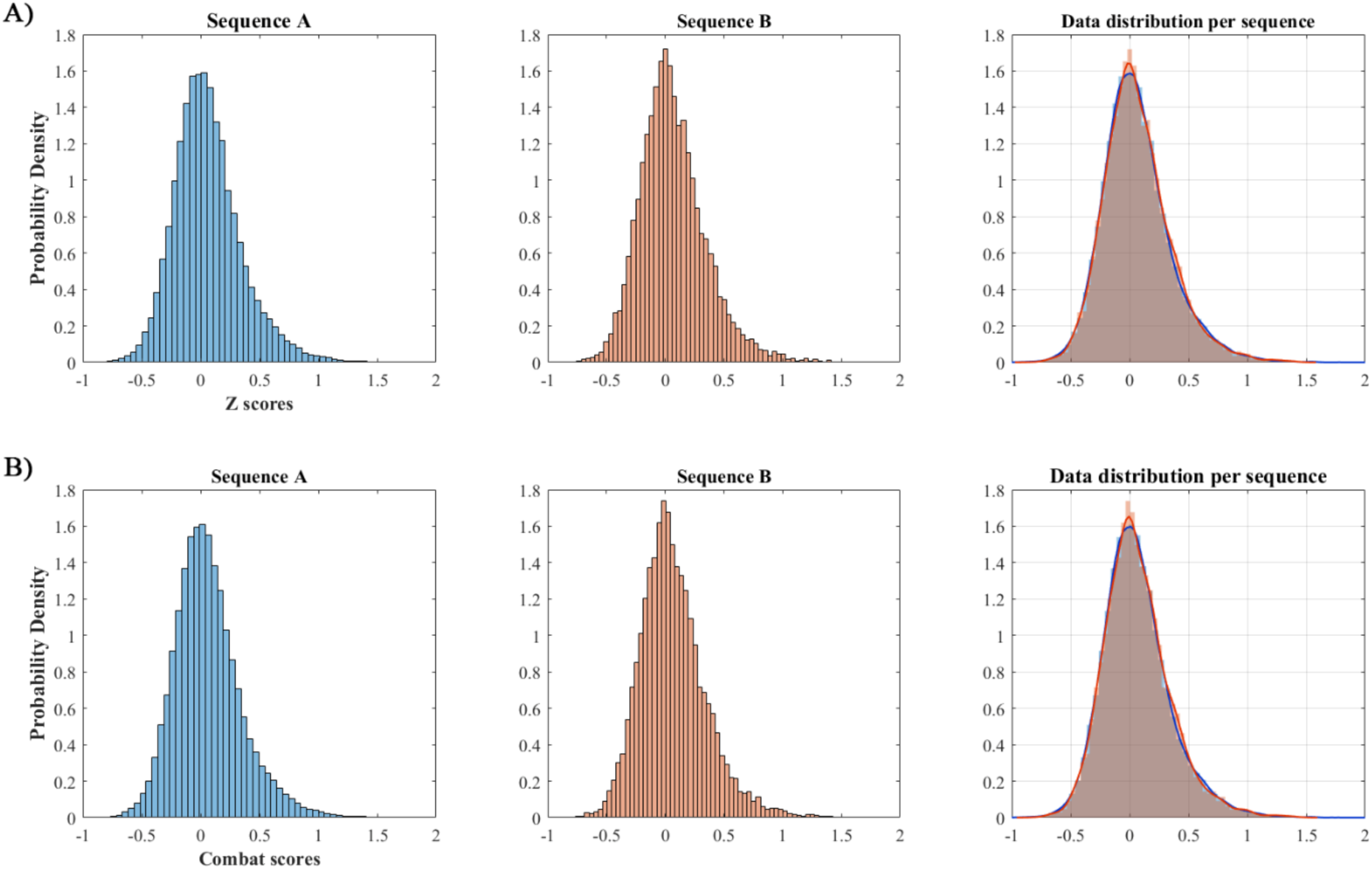
Sensitivity analysis of harmonisation methods for fMRI sequences. Sensitivity analyses comparing results before and after harmonisation using Z-scores and ComBat methods, which ensure homogeneous dynamic ranges across scanners. Results showed high concordance with initial analyses, with no significant differences between groups post-harmonisation.

## 3. Results

### 3.1. Pre-surgical functional connectivity patterns are altered compared to those of healthy participants

Pre-surgical rs-FC patterns in patients with left-hemisphere brain tumours differed significantly from those observed in healthy participants, as revealed by ROI-to-ROI analysis within the 8 pre-defined RSNs. Using the NBS non-parametric approach, which provides robust correction for multiple comparisons across connections, we identified specific bidirectional alterations. From the total of 496 connections examined among 32 ROIs, the NBS analysis revealed 8 connections with significantly altered functional strength between patients and healthy participants. Detailed information on these altered connections, including the specific RSNs involved, the direction of change (i.e., increased or decreased), and associated statistical metrics, is summarised in Table 3 (see also Figure 3). The reported differences correspond to effects that remain significant after applying cut-off thresholds designed to account for the substantial inter-individual variability inherent in a clinical sample, given that patients present lesions that vary in location, extent, type, and severity.

**Figure 3.**
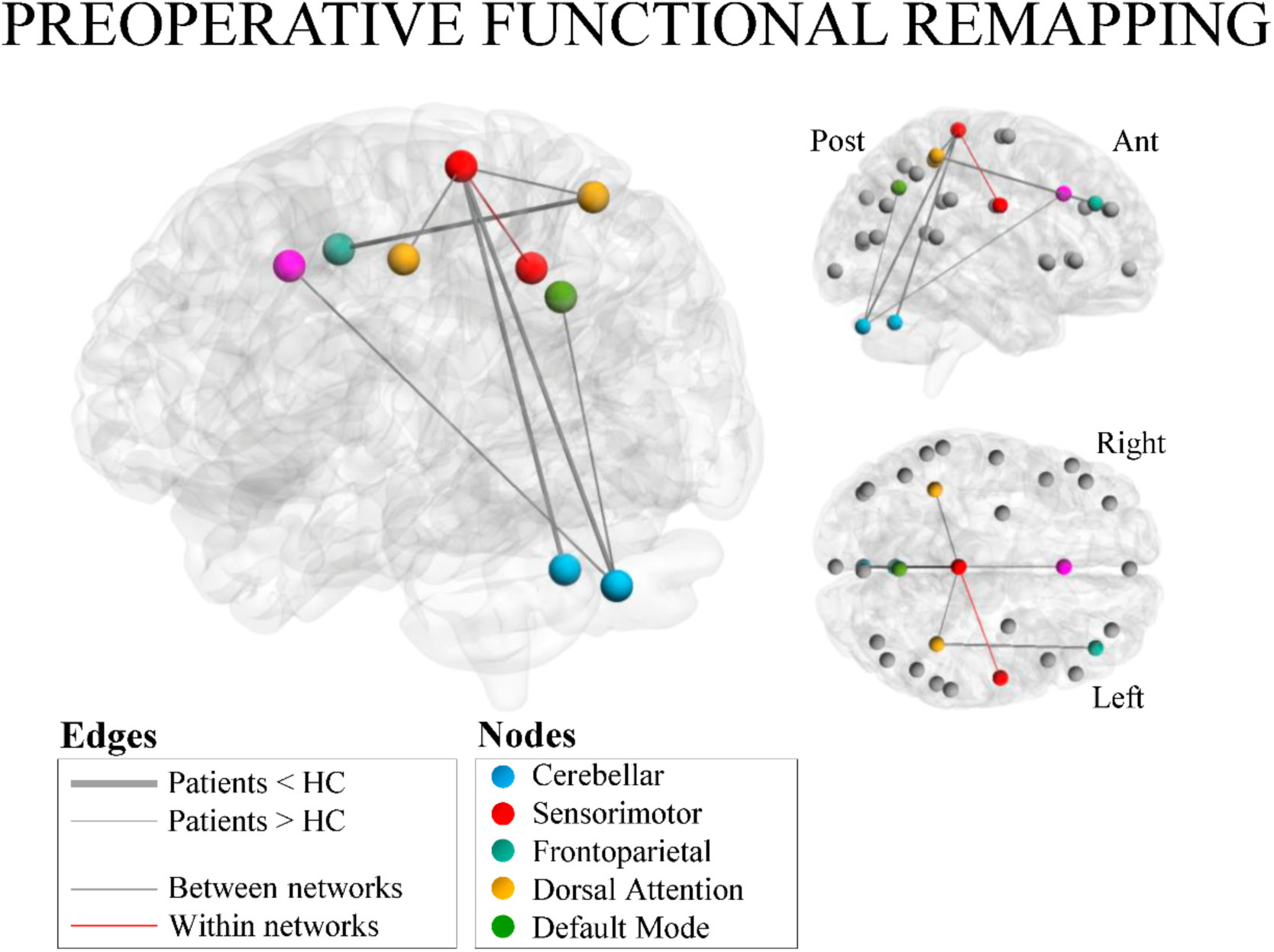
Functional connectome representing the results of the analysis comparing patients at the presurgical time point with healthy participants. Significant connections are displayed on a glass brain in a left sagittal view, slightly tilted to the right to allow full visualisation, including interhemispheric connections. Nodes within each network are shown in distinct colours. Thin edges indicate increased connectivity in patients relative to controls, whereas thick edges indicate decreased connectivity. On the right, two additional images (sagittal and axial) depict in grey all ROIs included in the analysis, along with the nodes involved in significant connections.

**Table 3.**
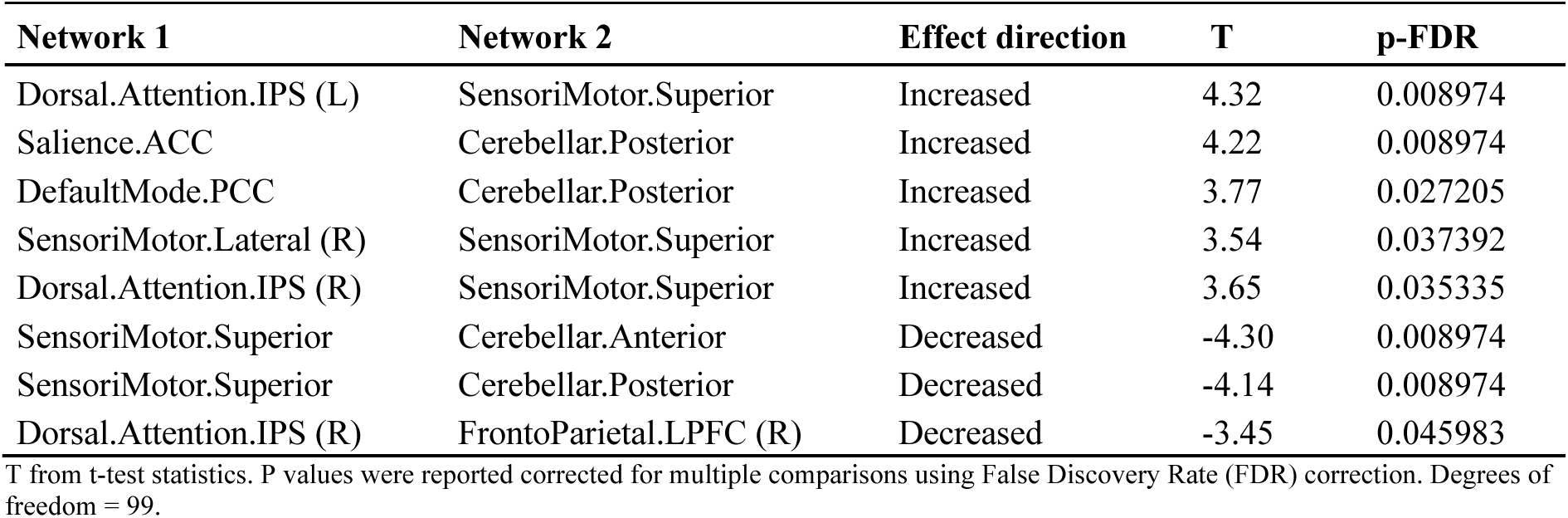
Results of the NBS analysis comparing patients and healthy participants in the presurgical stage.

In addition, the analysis based on graph properties revealed significant alterations in network segregation but not in network global efficiency before surgery. Specifically, patients showed reduced segregation relative to healthy controls. Although global efficiency did not differ significantly between groups, both metrics exhibited similar trends (see Figure 4).

**Figure 4.**
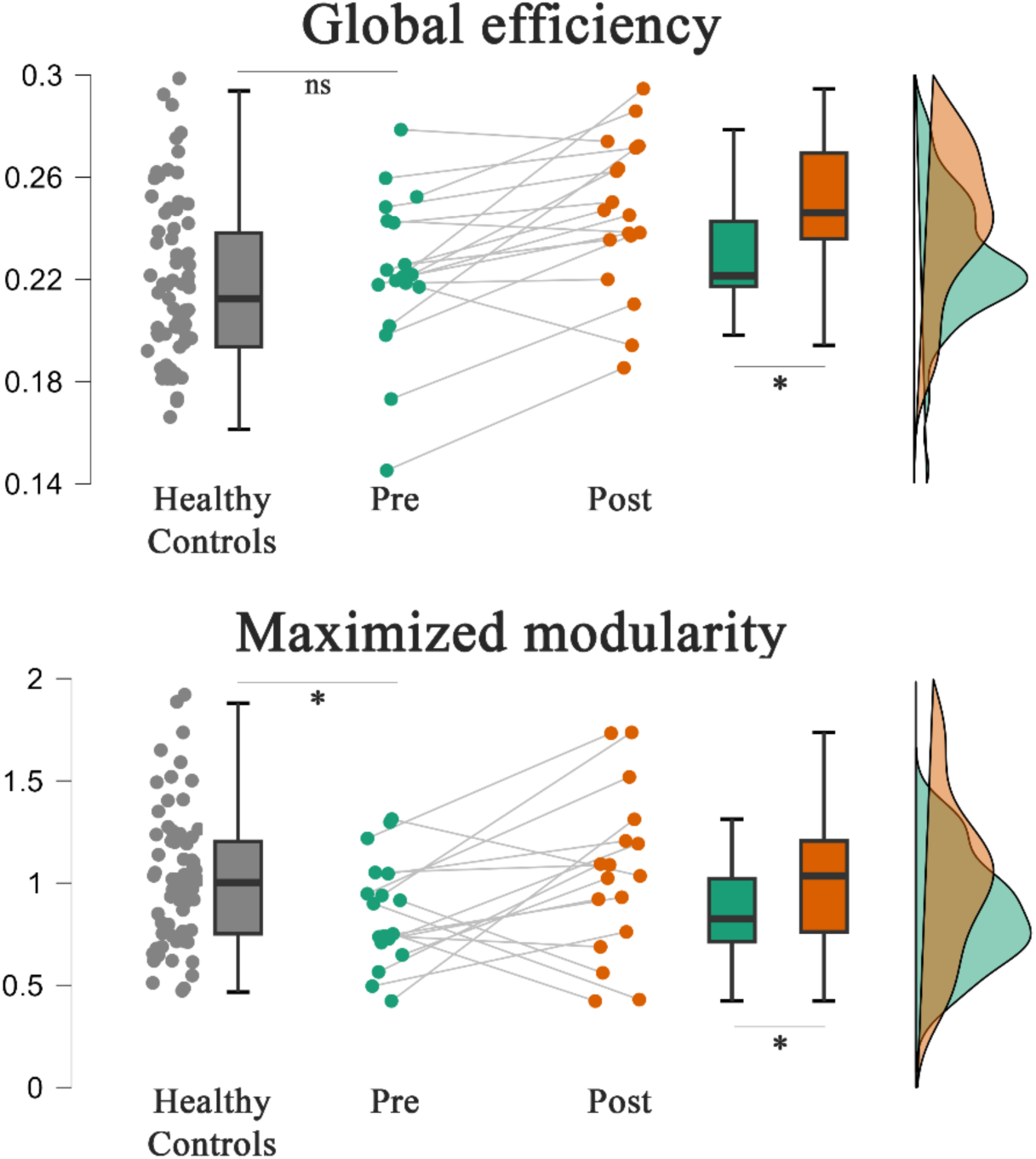
Global network efficiency and segregation. Comparisons of global efficiency (top) and maximised modularity, a measure of network segregation (bottom). Before surgery, patients showed significantly reduced modularity compared to healthy participants, while global efficiency was not significantly different. After surgery, patients exhibited a significant increase in both metrics compared to pre-operative levels.

### 3.2. Longitudinal changes between preoperative and postoperative stages

The surgical intervention preserved postoperative functional connectivity, and the differences relative to healthy controls remained stable over time. No significant longitudinal changes were detected between pre- and postoperative assessments across the 496 examined connections, based on the statistical thresholds described in the Methods section. However, the analysis based on graph properties revealed a significant postoperative increase in both network segregation and global efficiency compared to healthy controls (see Figure 4 and Table 4).

**Table 4.** Descriptive statistics for global efficiency and segregation, including pre- and postsurgical stages.

|  | **Global efficiency** |  |  | **Segregation** |  |  |
| --- | --- | --- | --- | --- | --- | --- |
|  | Healthy Participants | Pre | Post | Healthy Participants | Pre | Post |
| **Mean** | 0.221 | 0.223 | 0.246 | 1.002 | 0.857 | 1.092 |
| **Std. Deviation** | 0.031 | 0.031 | 0.030 | 0.331 | 0.260 | 0.448 |
| **Std. Error** | 0.004 | 0.007 | 0.007 | 0.040 | 0.061 | 0.106 |

### 3.3. Relationship between connectivity and behavioural variables

Individual cognitive progress was estimated based on behavioural measures, reflecting the difference in performance before and after surgery. These indices successfully captured the variability observed across patients, with some individuals showing cognitive preservation and others exhibiting signs of postoperative decline (see Figure 5). The cognitive prognosis index was then used as the outcome variable in the moderation analysis.

**Figure 5.**
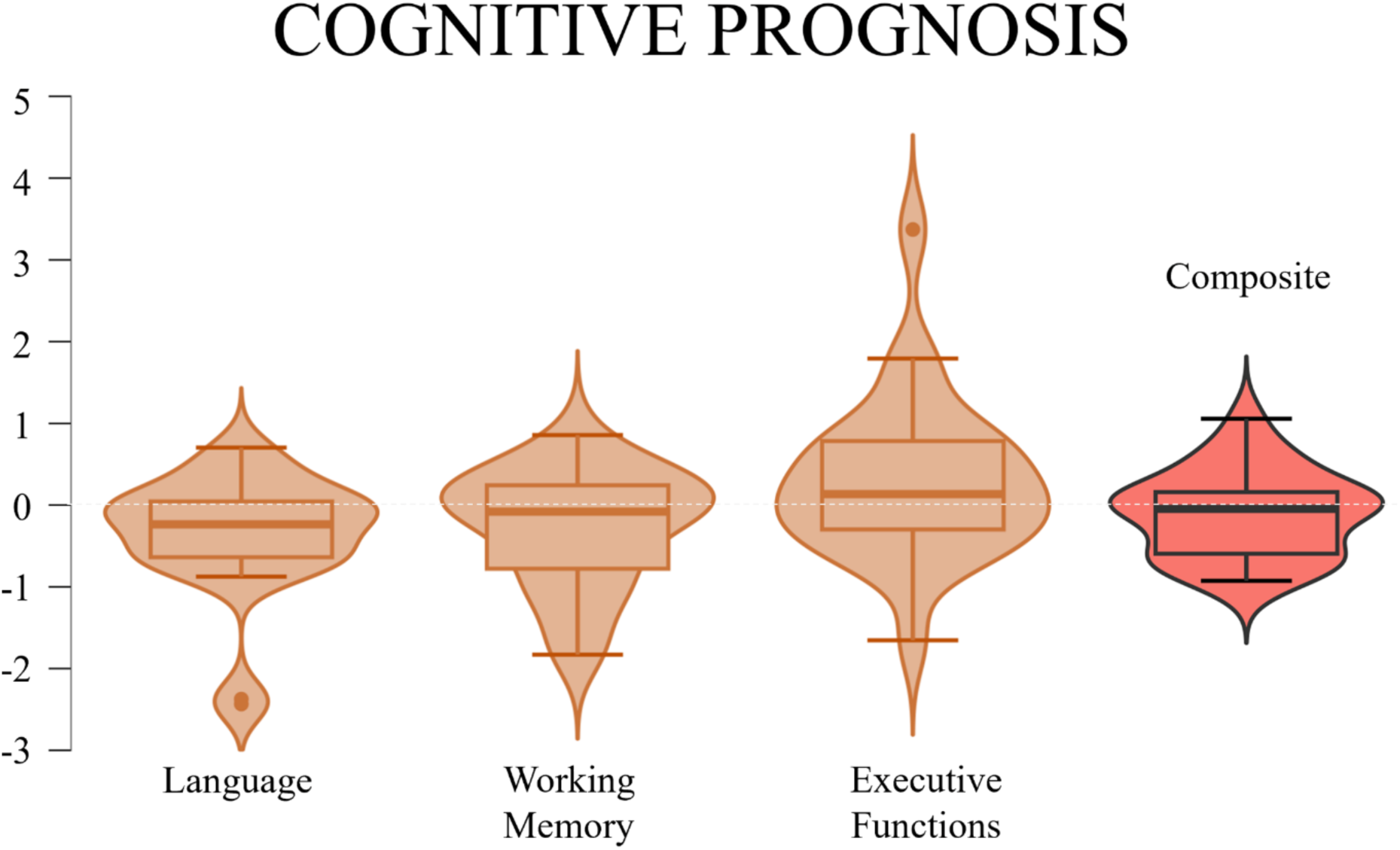
Means and standard deviations for each of the cognitive domains explored, corresponding to post- vs. preoperative differences. Note that both the composite scores by cognitive domain and the overall composite score successfully capture the effects observed in the patient sample, including individuals at both extremes of the distribution.

Global efficiency and network segregation were included as predictors in the moderation analysis, from which network segregation emerged as a predictor of cognitive prognosis. Patients with less segregated brain networks exhibited better cognitive recovery patterns. As shown in Figure 6, this direct effect was not moderated by clinical variables such as lesion volume or malignancy grade. However, although the moderating effect of age on the relationship between network segregation and cognitive improvement did not reach statistical significance (*p* = 0.266), a significant association was observed between age and cognitive progress (*p* = 0.006): younger patients showed more favourable cognitive recovery than older ones (see also Table 5).

**Figure 6.**
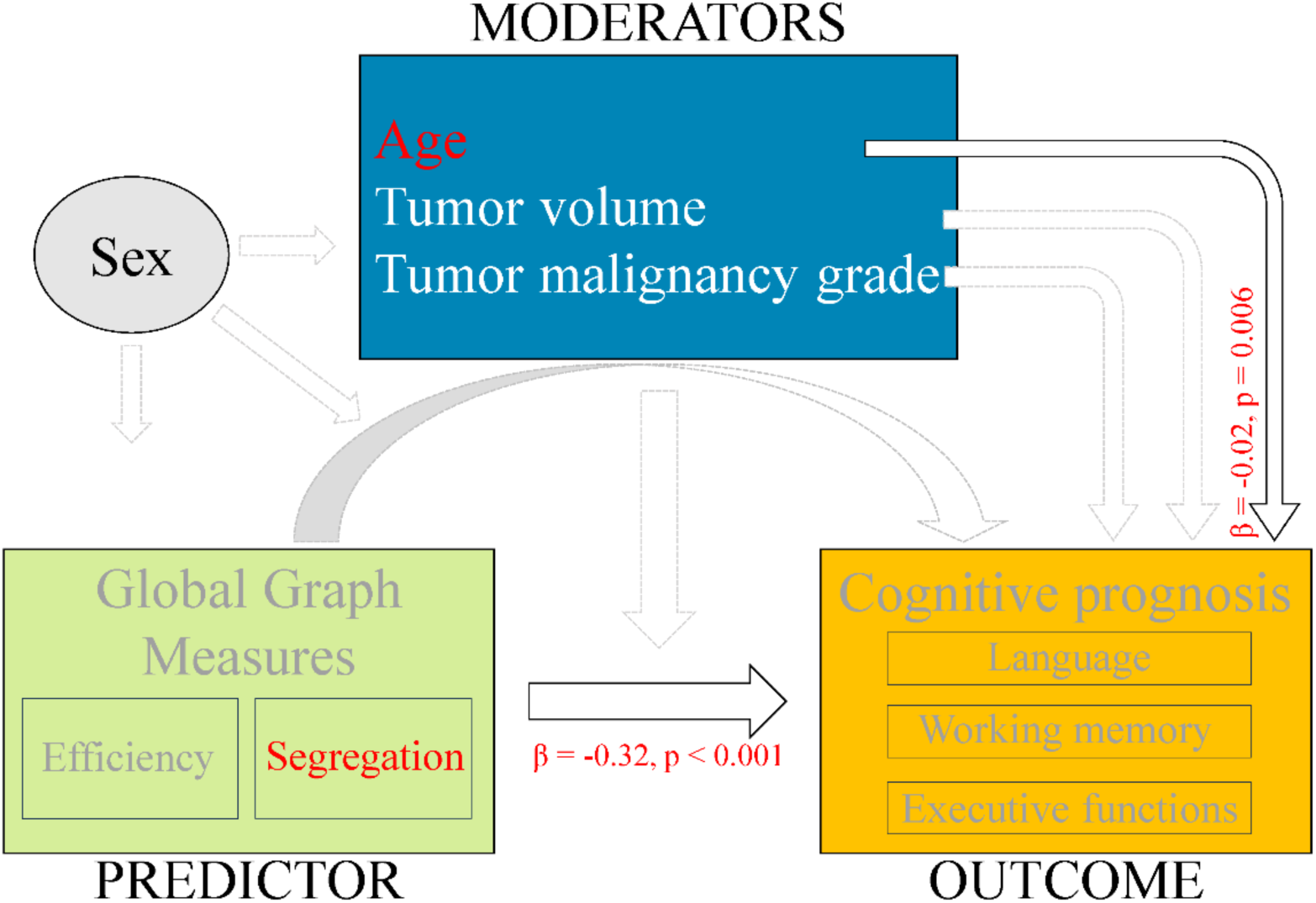
Diagram of the evaluated moderation model. Path coefficients and associated p-values are displayed along the arrows. Solid black arrows represent significant effects, whereas gray dashed arrows indicate non-significant effects.

**Table 5.** Moderation model results. Statistically significant values are indicated in bold within the p-value column and shaded in grey along the corresponding row.

| **Direct effects** | **Estimate** | **Std. error** | **Z value** | **p** |
| --- | --- | --- | --- | --- |
| Efficiency – Cog Progress | -1.688 | 1.661 | -1.016 | 0.310 |
| Modularity – Cog Progress | -0.322 | 0.073 | -4.421 | **< 0.001\*\*** |
| **Indirect effects** |  |  |  |  |
| Efficiency – Age – Cog Progress | -0.098 | 0.637 | -0.154 | 0.878 |
| Efficiency – Lesion volume – Cog Progress | 0.473 | 0.451 | 1.048 | 0.294 |
| Efficiency – Tumour Malignancy – Cog Progress | -0.341 | 0.418 | -0.817 | 0.414 |
| Modularity – Age – Cog Progress | -0.035 | 0.032 | -1.112 | 0.266 |
| Modularity – Lesion volume – Cog Progress | 0.047 | 0.039 | 1.211 | 0.226 |
| Modularity – Tumour Malignancy – Cog Progress | 0.019 | 0.023 | 0.817 | 0.414 |
| **Total effects** |  |  |  |  |
| Efficiency – Cog Progress | 0.937 | 0.054 | 17.30 |  |
| Modularity – Cog Progress | -0.296 | 0.084 | -3.506 | **< 0.001\*\*** |
| **Total indirect effects** |  |  |  |  |
| Efficiency – Cog Progress | 0.019 | 0.023 | 0.799 | 0.42 |
| Modularity – Cog Progress | 0.030 | 0.061 | 0.484 | 0.628 |
| **Path coefficients** |  |  |  |  |
| Age – Cog Progress | -0.020 | 0.007 | -2.745 | **0.006\*** |
| Lesion volume – Cog Progress | 0.166 | 0.160 | 1.042 | 0.298 |
| Tumour Malignancy – Cog Progress | -0.002 | 0.002 | -1.047 | 0.295 |

## 4. Discussion

In this study, we evaluated rs-FC changes in patients with left-hemispheric brain tumours across eight canonical networks. Four main findings were observed: 1) rs-FC is altered both inter- and intra-hemispherically in presurgical patients compared with healthy controls, providing evidence to consider brain tumors as a whole-brain disease; 2) these connectivity alterations remain largely stable four months after surgery, with efficiency levels comparable to those of healthy participants; 3) significant longitudinal changes in network topology were observed, indicating that both integration and segregation tend to recover towards a neurotypical pattern four months post-surgery; and 4) preoperative network segregation and age predicted cognitive recovery. Taken together, these results indicate that brain tumours induce network remapping, affecting functional connectivity of RSNs far beyond the regions strictly related to the affected region ^14,15^. The observed changes in network topological organisation likely reflect the adaptive role of domain-general networks in facilitating homeostatic processes.

We provide evidence that brain tumours, contrary to traditional views, induce functional changes beyond lesion location. Specifically, we identified changes in the connectivity patterns of the sensorimotor, salience, default mode, frontoparietal, and cerebellar networks, as well as among the nodes that constitute these networks. While alterations in those same networks have been previously documented for patients with brain tumours ^19,20,50^, their direction and potential implications as protective neuroplastic mechanisms remain unclear ^6^. We found those connections to occur bidirectionally: some connections were stronger while others were weaker in comparison to healthy participants. The existence of evidence pointing towards increased or decreased rs-FC in the same sample of patients reinforces the view that neuroplasticity encompasses a wide range of processes that go beyond any simplistic biological notions of “good” or “bad” ^51^, ultimately progressing towards balance. Moreover, as revealed by the mediation analysis, this pattern reflects an alteration in the efficiency of information flow, which may in turn stem from a reconfiguration of the neuronal hierarchy within domain-general networks. Analogous changes have been described in pathological ageing ^52^. Networks such as the DMN and the salience network adjust their responses to support specific functions that, in the healthy brain, are primarily sustained by specialised networks. In this context, our findings align with previous observations of altered activation in domain-general networks during task performance in tumour patients. For instance, using the same sample as in the present study, increased recruitment of the DMN during language production tasks has been reported and interpreted as a compensatory mechanism ^24^. Tumour invasion of neural tissue sustaining language functions not only induces a functional remapping of the language network but also extends to resting state domain-general networks, as a compensatory mechanism that reshapes the synchronised activity across subnetworks.

We found bilateral increased connectivity between the dorsal attention and superior sensorimotor networks. Similar changes between the dorsal attentional network and the FPN have been documented ^12^, along with increased connectivity within the DMN and between the DMN and other brain networks ^53^, with the DMN being proposed to participate in supporting cognition after surgery ^54^. Decreases in rs-FC were observed between the dorsal attention IPS and the frontoparietal LPFC in the right hemisphere. In the literature, changes regarding the FPN are ambiguous, with increased rs-FC in some studies ^12^ and globally decreased strength in others ^55^. Significant decreases in FC have been shown to indicate a disruption of network interactions with the potential to affect behaviour, as seen in previous studies ^13^; however, we did not find those associations. The increase in cerebro-cerebellar connections, already present at the presurgical stage, could be seen as a supporting mechanism promoting adaptive responses for language ^56^.

A promising hypothesis, which interprets the configuration of RSNs as an adaptive compensatory mechanism, suggests that the DMN may lead to the recruitment of specific underlying networks for a given task, controlling the hierarchy of neural network dynamics ^57^. However, the role of specific networks within this hierarchy remains an open question. Our results extend this inquiry to dynamic changes among domain-general networks, as changes were not limited to the DMN and included networks with comparable contributions to overall functional organisation ^58^. The recruitment of domain-general networks holds the potential to facilitate efficient responses in a demanding and dynamic environment, preserving homeostasis. There is still the possibility that a sustained involvement of these networks might lead to neuronal overload and, eventually, may become detrimental for cognitive recovery. However, according to our results, the involvement of domain-general networks is associated with improvements in cognitive performance. Further, network efficiency during the presurgical stage could be used as a predictor of how cognitive performance would evolve.

Unlike previous studies, the functional connectivity changes observed during the presurgical phase in our sample remained stable four months after surgery when comparing connectivity matrices comprising 496 connections ^18,21^. Although this analysis did not reveal significant longitudinal changes, likely due to a combination of limited sample size and substantial inter-individual variability, global graph properties did capture postsurgical changes, as described in previous reports ^3,25,26,54^. Specifically, both segregation and integration metrics initially diverged further from control values and showed a trend to resemble the neurotypical pattern after surgery.

The observed changes in network integration and segregation towards a neurotypical pattern after surgery support the idea that when surgical intervention succeeds in preserving the neuroplastic changes that occurred in the brain prior to surgery, adaptive mechanisms continue to facilitate postoperative recovery. Consistently, previous studies have shown that glioma patients exhibit an altered global network organisation at rest, with changes in graph metrics such as clustering coefficient and path length, which were found to correlate with patients’ cognitive status ^25,26,28–30^. These alterations revealed reduced network efficiency and a more random topology in glioma patients compared with healthy controls. Diffusion imaging studies in glioma patients have also reported that preoperative structural network differences relative to controls tend to diminish after surgery ^4,55^. These findings are in line with the longitudinal functional changes demonstrated in the present study, suggesting a disruption of the optimal homeostatic balance followed by recovery of the baseline state after surgical tumour removal.

Discrepancies among previous studies reporting significant temporal changes in neuroplasticity mechanisms—particularly those comparing connectivity matrices as dependent variables in tumour populations undergoing similar surgical approaches—should be interpreted with caution. In several of these studies, oncological interventions were combined with cognitive rehabilitation programs designed to optimise language function recovery ^17,18^. A direct comparison between our patients and those examined in these works would be valuable to determine the extent to which rehabilitation techniques can reactivate or enhance spontaneous plasticity mechanisms. However, the postoperative changes described in such studies may primarily reflect the effects of cognitive intervention rather than the spontaneous neuroplastic processes specifically addressed in the present work. Conversely, other published studies include patients with glioblastoma—a markedly more aggressive pathology—where the observed functional changes are likely driven by accelerated cognitive decline and rapid disease progression ^21^.

Although direct evidence is still needed to demonstrate the capacity of neurons to form new synapses or strengthen alternative pathways in oncological conditions, the present study suggests that the brain’s intrinsic plasticity mechanisms remain preserved despite the progression of the lesion. The degree of functional network recovery, considering the relationship demonstrated in this study between age and cognitive recovery, varies across individuals. Age is a crucial determinant of the brain’s neuroplastic potential, directly influencing the process of postoperative cognitive recovery ^59,60^.

In summary, our findings prove that the effects of brain tumours are not strictly localised phenomena but spread across the brain. This perspective aligns with considering gliomas as a whole-brain systemic disease ^61^, with both inter- and intra-hemispheric alterations ^14,62^. The bidirectional nature of the connectivity changes observed in this study reflects the brain’s dynamic capacity for adaptation that is not confined to increases in activations. Our results also support the promise of incorporating graph-theory measures to study topological properties of the networks in patients. Finally, the role of the cerebellum in supporting cognitive functions requires further attention, as its involvement may be crucial for long-term recovery and compensation.

### 4.1. Limitations

While providing valuable insights, our findings should be interpreted in light of certain limitations. Despite the difficulties inherent in collecting patient data, the sample size in this study is still limited, particularly for the longitudinal follow-up. This fact restricted our ability to correlate cognitive performance with rs-FC to draw strong conclusions. Consistently linking rs-FC metrics with cognitive performance using a larger sample could provide clarity as to whether they represent compensation and strengthen their effectiveness as a predictor to guide therapeutic decisions ^63^. Further, in future work, both behavioural and rs-FC data should be combined with structural data to obtain a more comprehensive picture.

Regarding the longitudinal evaluation, although fixed-time assessments are highly valuable, it remains uncertain whether this approach captures stable indicators for optimal recovery. Recovery seems to depend largely on surgical intervention practices; therefore, understanding its relationship with rs-FC changes would be beneficial. Future research should work on establishing guidelines to include optimal re-scan intervals ^6^, which might vary depending on tumour grade and its associated temporal progression. Mechanisms such as abnormal vascular architecture ^64^ and blood flow characteristics ^65^ can modulate the BOLD signal within tumour regions, influencing FC outcomes across RSNs. Neurovascular uncoupling, resulting from disruptions in neuron-astrocyte interactions that regulate cerebral blood flow, further complicates rs-FC interpretation ^66^. There is a growing need for tools that can better elucidate the modulation of BOLD fMRI signals in patients with brain tumours, thereby improving both the reliability and interpretability of functional connectivity findings. Cerebrovascular reactivity has emerged as a promising approach to provide physiological insight into these signal changes ^67,68^.

Another limitation is that in whole-brain RSNs studies, the choice of parcellation scheme significantly impacts the results, influencing how we interpret rs-FC across the brain. These schemes vary widely in their resolution and anatomical definitions, each introducing specific challenges. In our study, we used the CONN toolbox to apply a data-driven parcellation derived from a large normative sample. This approach offers strong anatomical interpretability and holds promise for clinical translation, as it enables standardisation and reduces reliance on the subjective expertise of clinical staff. Nevertheless, the relatively coarse resolution of this parcellation constrains the ability to detect differences that may occur within neural networks at a finer spatial scale. In future studies, voxel-level methods could also be explored, as they could provide finer-grained insights ^69^. Finally, to improve the clinical applicability of resting-state data, moving towards single-subject evaluations of rs-FC could be of great interest. More individualised clinical insights could help integrate resting-state data into routine clinical practice ^70,71^.

## Conclusions

This study demonstrates that brain tumours have a widespread impact on RSNs, supporting the conceptualisation of brain tumours as a whole-brain disease. At the network level, tumour-related alterations affected both local and global topological organisation preoperatively: patients exhibited decreased segregation while maintaining global efficiency relative to healthy controls, along with changes in the synchronisation of domain-general networks. Moreover, significant longitudinal changes were observed between the pre- and postoperative phases, characterised by postoperative increases in both segregation and integration estimates that approximated the neurotypical pattern. Preoperative network segregation was found to be a significant predictor of postoperative cognitive recovery, with potential as a marker of resilience. This perspective underscores the need to reconsider compensatory mechanisms and refine strategies for targeted therapeutic intervention to adjust cognitive recovery expectations. Future investigations involving larger and more diverse cohorts, together with advanced network modelling, are needed to clarify the mechanisms of neural plasticity underlying long-term cognitive and functional outcomes in patients with brain tumours.

## Acknowledgments

The authors would like to thank all the participants who agreed to take part in this study and the lab team for assisting with data collection, especially David Carcedo and Maite Kaltzakorta.

## Data Availability

The data are not publicly available due to the data-sharing policies of the different institutions involved concerning vulnerable clinical information.

## Authorship

Conceptualization, L.M.-O., I.Q., M.C., I.D.; Methodology, L.M.-O., I.Q., I.D.; Software, L.M.-O., I.Q., I.D.; Validation, I.Q.; Formal analysis, L.M.-O., I.D., I.Q.; Investigation, L.M.- O., I.Q.; Resources, L.A., G.B., S.G.-R., I.P., M.C., I.Q., Writing—original draft preparation, L.M.-O., I.D., I.Q.; Writing—review and editing, all authors; Visualization, L.M.-O., I.Q.; Supervision, I.Q., M.C., I.D.; Project administration, L.A., I.Q.; Funding acquisition, L.A., M.C., I.Q. All authors have read and agreed to the published version of the manuscript.

## Informed Consent Statement

Informed consent was obtained from all subjects involved in this study.

## Supplementary Material

**Table 1S.** Non-missing values for each measure at each time point.

| **Cognitive Domain** | **Measure** | **Pre**<br>**(n = 35)** | **Post**<br>**(n = 24)** |
| --- | --- | --- | --- |
| **Language** | Language production task | 34 | 22 |
|  | Pseudoword reading task | 25 | 19 |
|  | Reading comprehension task | 25 | 18 |
|  | Sentence picture matching task | 24 | 19 |
|  | Grammatical judgment task | 25 | 19 |
|  | Verbal intelligence task | 35 | 24 |
| **Working Memory** | Forward and backward digit span | 31 | 20 |
|  | Forward digit/letter span | 31 | 20 |
| **Executive functions** | Non-verbal intelligence task | 26 | 24 |
|  | Cross-out | 27 | 22 |
| **Composite variable** | Language | 37 | 24 |
|  | Working memory | 31 | 20 |
|  | Executive Functions | 27 | 24 |
|  | Global | 35 | 24 |

**Table 2S.** Resting-state fMRI parameters for the two functional imaging sequences.

|  | **Sequence A** | **Sequence B** |
| --- | --- | --- |
| Voxel resolution (mm) | 3 mm<sup>3</sup> | 2 mm<sup>3</sup> |
| Repetition time (ms) | 2000 ms | 850 ms |
| Echo time (ms) | 30 ms | 35 ms |
| Image columns | 384 | 792 |
| Image rows | 384 | 792 |
| Flip Angle (°) | 78° | 56° |
| Slices | 32 | 66 |
| Matrix Size | 91 x 109 | 91 x 109 |
| Acceleration factor | 0 | 0 |
| Echo Spacing (ms) | 64 ms | 88 ms |
| Echo planar images | 180 | 1058 |

